# Matthew effect: common species become more common and rare ones become more rare in response to artificial light at night

**DOI:** 10.1101/2021.08.09.455752

**Authors:** Yanjie Liu, Benedikt Speißer, Eva Knop, Mark van Kleunen

**Affiliations:** Key Laboratory of Wetland Ecology and Environment, Northeast Institute of Geography and Agroecology, Chinese Academy of Sciences, Changchun 130102, China; Ecology, Department of Biology, University of Konstanz, 78464 Konstanz, Germany; Agroscope, Agroecology and Environment, Zürich, Switzerland; Department of Evolutionary Biology and Environmental Studies, University of Zürich, Zürich, Switzerland; Zhejiang Provincial Key Laboratory of Plant Evolutionary Ecology and Conservation, Taizhou University, Taizhou 318000, China

**Keywords:** Anthropocene, exotic, invasiveness, light pollution, non-native, plant-insect interaction, trophic level

## Abstract

Artificial light at night (ALAN) has been and still is rapidly spreading, and has become an important component of global change. Although numerous studies have tested its potential biological and ecological impacts on animals, fewer have tested its impacts on plants, and very few studies have tested whether it affects alien and native plants differently. Furthermore, common plant species, and particularly common alien species, are often found to benefit more from additional resources than rare native and rare alien species. Whether this is also the case with regard to increasing light due to ALAN is still unknown.

Here, we tested how ALAN affects the performance of common and rare alien and native plants directly and indirectly via flying insects. We grew five common alien, six rare alien, five common native and four rare native plant species under four combinations of two ALAN (no ALAN *vs* ALAN) and two insect-exclusion (no exclusion *vs* exclusion) treatments, and compared their biomass production.

We found that common plant species, irrespective of whether they are alien or native, produced significantly more biomass than rare species, particularly under ALAN. Furthermore, alien species tended to show a slightly stronger positive response to ALAN than native species (marginally significant origin × ALAN interaction, *p* = 0.079).

Our study shows that common plant species benefited more from ALAN than rare ones. This might lead to shifts in plant diversity and vegetation composition, further propelling global biodiversity decline, when ALAN becomes more widespread. In addition, the slightly more positive response of alien species indicates that ALAN might increase the risk of alien plant invasions.

## Introduction

Light pollution due to artificial light at night (ALAN) has increased dramatically in the last century, and has altered the natural night-time environment in large areas of the Earth ^1,2^. As a consequence, ecological impacts of ALAN have become an important focus for global change research in recent years ^3-9^. To date, most research on the impact of ALAN has focused on the behavior, physiology, and life history of animals ^10-13^, and only a few studies have focused on plants ^6,12,14^.

ALAN might interfere with the plants’ seasonal timing, and thereby affect plant phenology ^6^. In many cases, ALAN (e.g. street lighting) is sufficiently bright to induce a photosynthetic response ^15^, and thus could directly influence growth and resource allocation of plants ^6,15^. Given that such direct impacts generally vary among plant species ^12,15,16^, ALAN is likely to affect the composition of the vegetation ^16^. Therefore, it is not unlikely that ALAN may also influence alien and native plants differently, and consequently affect the process of alien plant invasion. For example, Murphy et al. ^17^ did field surveys in urban areas, and found that the presence of the invasive plant *Bromus tectorum* was positively associated with the presence of streetlights. To the best of our knowledge, only one previous empirical study has tested how ALAN could affect the performance of alien and native plants. In a competition experiment between alien and native species, Speißer et al. ^15^ found a trend that naturalized alien plants took less advantage of ALAN than less-widely naturalized alien plants. This could imply a turn-over in invasive alien species when ALAN continues to increase.

Numerous empirical studies have tested how alien and native plants respond to resource increases, and found mixed results ^18-21^. This may partly depend on how widespread or dominant the alien and native plants are that were compared ^18,20,22-24^. For example, Dawson et al. ^18^ found that common species take more advantage of nutrient addition than rare species, whether they are alien or native. In other words, common alien plants do not necessary benefit more from resource increase than common native plants ^20^. Furthermore, common species are often considered to have traits conferring greater light capture ability, such as a high specific leaf area, and consequently they may respond more positively to light addition than rare species ^25,26^. Given that ALAN is an increase in the resource light, it is plausible that common species might benefit more from ALAN than rare species. Therefore, rigorous tests of whether ALAN might affect future plant invasions, should also consider how common or rare the alien and native species are ^15,18,24,27^.

ALAN can also affect other organisms such as insects ^9,28-30^, which could play very important roles, e.g. as herbivores and pollinators, in regulating plant growth and reproduction. Consequently, ALAN could indirectly affect plants by modifying plant-insect interactions ^7,9,28,31-33^. Furthermore, alien plants, due to their evolutionary novelty, might attract different numbers and types of herbivores ^34-36^ and pollinators ^37-39^ than native plants. Following this logic, ALAN might have different indirect effects, mediated by other trophic levels, on the performances of alien and native plants. To the best of our knowledge, however, this idea has never been tested.

To test direct and indirect effects of ALAN on the performance of common and rare aliens and natives, we conducted a multispecies, common-garden experiment. We compare the biomass production in response to ALAN with and without insect-exclusion treatments among five common alien, six rare alien, five common native and four rare native plant species. We address the following specific questions: 1) Does ALAN directly affect biomass production of plants, and does it matter whether the species is common or rare and alien or native? 2) Does ALAN affect biomass production of plants indirectly via effects on insects, and does it matter whether the species is common or rare and alien or native?

## Material and Methods

### Study Species

To investigate effects of ALAN on common alien, rare alien, common native and rare native species, we selected a total of 20 terrestrial grasses and forbs co-occurring in grasslands of Germany. To cover a wide taxonomic breadth, these species were chosen from six different families. To prevent possible taxonomic bias regarding origin and commonness, we tried to include for each family at least one species for each of the four categories. However, due to poor germination and limited seed availability for some species, our final species set was not fully balanced with regard to taxonomy; we used five common alien, six rare alien, five common native, and four rare native species (Table S1). We classified the species as naturalized alien or native to Germany based on the BiolFlor database (www.ufz.de/biolflor). As species commonness has multiple dimensions ^40,41^, we classified the species as common or rare based on both occupancy frequency and local abundance level (Table S1). Specifically, we assigned a species as common if it is locally abundant (i.e. can form large groups) and occupies more than 900 out of all 3000 grid cells in Germany, and as rare if it is not locally abundant and occupies fewer than 400 grid cells, according to the FloraWeb database. Seeds of the species used in the experiment originated from botanical gardens, commercial seed companies, or from wild populations (Table S1).

### Experimental ALAN and Insect-Exclusion Facility

To simulate the ALAN and insect-exclusion treatments, we arranged 20 metal cages (2 m × 2 m × 2 m) on an 21 m × 8 m area (Fig. 1) at the Botanical Garden of the University of Konstanz, Germany (N: 47°69’19.56”,E: 9°17’78.42”,). We assigned each of the 20 cages to one of the four combinations of two ALAN (no ALAN *vs* ALAN) and two insect-exclusion (no exclusion *vs* exclusion) treatments. In other words, each of the four treatment combination had five replicate cages. To impose ALAN, we randomly selected 10 of the cages, and installed LED spotlights (LED-Strahler Flare 10 W, IP 65, 900 lm, cool white 6500 K; REV Ritter GmbH, Mömbris, Germany), which were switched on each day from sunset to sunrise. Activity of the lamps was controlled by a photoelectric switch (DÄ 565 08, Eberle Controls GmbH, Nürnberg, Germany), turning the lamps on at an ambient light intensity below 30 lux, and switching them off if ambient light intensity exceeded 30 lux. To reduce lateral light radiation of the LED spotlights, we used plastic boxes (40 cm × 50 cm × 27 cm) as lampshades. The LED spotlights, which emit photosynthetically active radiation (Fig. S1), were fixed at 2 m height in the center of each cage (Fig. 1). To achieve a realistic light intensity, as can be found under street lights, two layers of white tissue were placed beneath the spotlights. This way, the final light intensity was 23.8 ± 1.2 lux, which is within the range of light intensities at ground level under street lights ^6^. The remaining ten cages, serving as ambient light treatments, also had the lampshades installed, but without LED spotlights. Light intensity at floor level in the cages without ALAN was 0.06 ± 0.06 lux.

**Figure 1.**
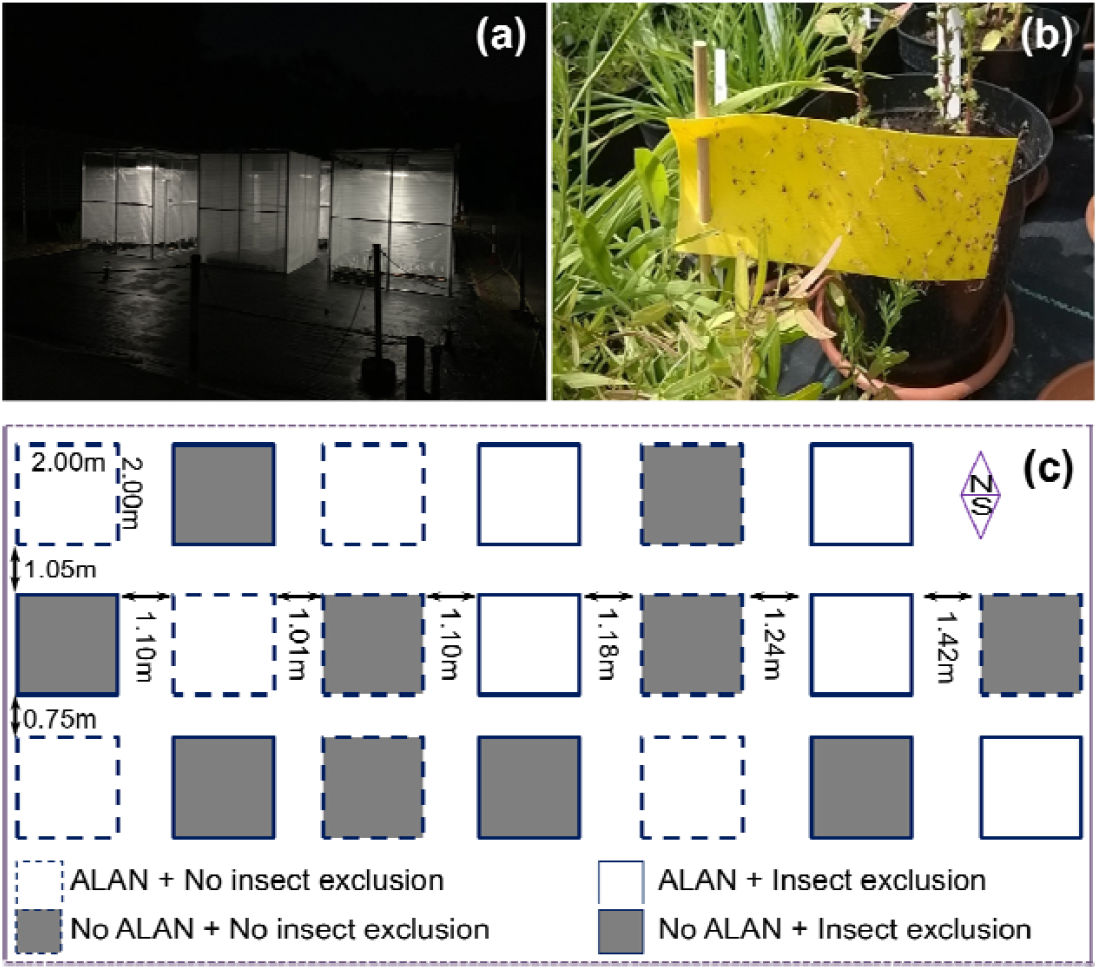
Illustration of the experimental setup. (a) Photograph of the experiment at night. (b) Photograph of the insect traps within the cages to monitor insect abundance for the different treatments. (c) The locations of the cages and their assignment to the four combinations of two ALAN and two insect-exclusion treatments outside in the Botanical Garden of the University of Konstanz.

To impose the insect-exclusion treatment, we randomly assigned five of the cages with and five of the cases without ALAN to the insect-exclusion treatment. The sides and roofs of all 20 cages were covered with a metal grid with a mesh size of 3 cm × 5 cm, allowing insects to enter. The ten insect-exclusion cages, we completely covered (sides and roof) with insect net (mesh size: 0.4 mm × 0.77 mm, 115 g / m^2^; FVG Folien-Vertiebs GmbH, Dernbach, Germany). To minimize differences in light intensity and rain shelter effects between the treatments with and without insect-exclusion, we also partly covered the cages that were not part of the insect-exclusion treatment with insect net. The roofs of these cages were completely covered, while at each side a 1.5 m wide strip was attached. This way 25-cm wide strips at the bottom and top of each side were not covered, and allowed insects to enter the cages. To test the effectiveness of the insect-exclusion treatment, we implemented during three days two sticky, yellow insect traps in each cage (see Fig. 1b). Counts of the caught insects showed that the number of flying insects was 70.6% lower in the insect-exclusion cages than in the control cages, and that it was 717.4 % higher in the cages with ALAN than in the cages without ALAN (Table S2; Fig. S2).

### Precultivation and Experimental Setup

On 13 May 2019, we started to sow the study species separately into trays (13.4 cm × 12.2 cm × 4.9 cm) filled with potting soil (Einheitserde, Pikiererde CL P). To obtain seedlings of a similar developmental stage at the start of the experiment, we sowed the species on different dates (Table S1), based on germination experience with those species from previous studies. After sowing, we kept all trays in a greenhouse with a temperature between 18 and 21°C, and a day:night cycle of approximately 16:8 hr. On 3 and 4 June 2019, we transplanted similar-sized seedlings into 2-L circular plastic pots filled with the same type of potting soil as used for germination. In each pot, we planted three individuals of the same species. To maximize the use of germinated seedlings and increase the statistical power of the study, we transplanted 33 pots for each species, resulting in total of 693 pots (i.e. 99 seedlings per species and 2,079 seedlings in total).

After transplanting the seedlings, we distributed the 693 pots over the 20 cages with the four combinations of ALAN × insect-exclusion treatments. We replaced dead seedlings on 9 and 10 June 2019. For each species, we first distributed 20 of the 33 pots over the 20 cages, and then randomly assigned the remaining 13 pots to 13 of the 20 cages. In other words, each cage had one to two pots for each of the 21 species. To distribute the pots as evenly as possible among the cages, we had 35 pots for each of 13 cages, and 34 pots for each of the remaining seven cages. Within each cage, the pots were randomly assigned to fixed positions and were re-randomized every 14 days. To prevent nutrient limitation during the experiment, we fertilized all pots weekly (with 1 ‰ [w/v] Universol^®^ blue oxide, ICL SF Germany & Austria, Nordhorn, Germany) from 4 June 2019 (i.e. four weeks after the start of the experiment) onward. We also watered the plants regularly to keep the substrate moist throughout the entire experiment.

From 13 to 15 August 2019, ten weeks after transplanting, we harvested the plants. As the roots could not be extracted from the potting soil, we only harvested the aboveground biomass of each pot. All aboveground biomass of each pot was dried at 70°C for at least 72 hours, and then weighed.

### Statistical analysis

To analyze the effects of ALAN, insect exclusion and their interactions on the performance of common alien, rare alien, common native and rare native species, we fitted linear mixed-effects model using the lme function of the ‘nlme’ package ^42^ in R 4.0.3 ^43^. Aboveground biomass production of each pot was the response variable. To meet the assumption of normality, aboveground biomass production was square-root-transformed. We included ALAN treatment (no ALAN *vs* ALAN), insect-exclusion treatment (no exclusion *vs* exclusion), species origin (alien *vs* native), species commonness (common *vs* rare), and all their interactions as fixed effects in the model. To account for non-independence of individuals of the same species and for phylogenetic non-independence of the species, we included species identity nested within family as random effects in the model. In addition, to account for non-independence of plants that were in the same cage, we also included cage as a random effect in the model. As the homoscedasticity assumption of the model was violated, we included variance structures to model different variances per species using the “*varIdent*” function in the R package ‘nlme’ ^42^. We used log-likelihood ratio tests to assess significance of the fixed effects ALAN treatment, insect-exclusion treatment, species origin, species commonness, and their interactions ^44^.

## Results

Averaged across all treatments and origins, the common species produced significantly more aboveground biomass (mean: 6.6 g) than the rare species (3.9 g; Table 1; Fig. 2 and Fig. 3). Whereas common species responded positively to ALAN (+9.2%), the reverse was true for rare species (−2.0%; significant C × L interaction in Table 1; Fig. 3). Although we did not find any significant main effects for species origin and ALAN treatment, alien species tended to respond more positively (+7.7%) to ALAN than native species (+2.3%; Fig. 3). This was reflected in a marginally significant O × L interaction (Table 1). We found no significant main effect of the insect-exclusion treatment, and no significant interactions of insect exclusion with ALAN, commonness or origin (Table 1).

**Table 1.** Results of linear mixed-effect model testing the effects of species origin (native *vs* alien), commonness (common *vs* rare), ALAN treatment (no ALAN *vs* ALAN), insect-exclusion treatment (no exclusion *vs* exclusion) and their interactions on aboveground biomass production. Significant effects (*P* < 0.05) are in bold and marginally significant effects (*P* < 0.1) are underlined.

| **Fixed effects** | Biomass (sqrt-transformed) |  |  |
| --- | --- | --- | --- |
| | df | $\chi^2$ | P |
| Origin (O) | 1 | 1.323 | 0.250 |
| Commonness (C) | 1 | 5.155 | **0.023** |
| ALAN (L) | 1 | 0.191 | 0.662 |
| Insect (I) | 1 | 1.207 | 0.272 |
| O × C | 1 | 1.215 | 0.271 |
| O × L | 1 | 3.095 | <u>0.079</u> |
| O × I | 1 | 0.210 | 0.647 |
| C × L | 1 | 6.779 | **0.009** |
| C × I | 1 | 0.297 | 0.586 |
| L × I | 1 | 0.009 | 0.927 |
| C × L × I | 1 | 0.054 | 0.817 |
| O × L × I | 1 | 0.428 | 0.513 |
| O × C × I | 1 | 0.609 | 0.435 |
| O × C × L | 1 | 0.811 | 0.368 |
| O × C × L × I | 1 | 0.352 | 0.553 |
| **Random effects** |  | **SD** |  |
| Family |  | 0.251 |  |
| Species* |  | 0.469 |  |
| Cage |  | 0.097 |  |
| Residual |  | 0.437 |  |
| **R<sup>2</sup> of the model** | **Marginal** | **Conditional** |  |
|  | 0.163 | 0.670 |  |

\*Standard deviations for individual species random effects for the saturated model are found in Table S3.

**Figure 2.**
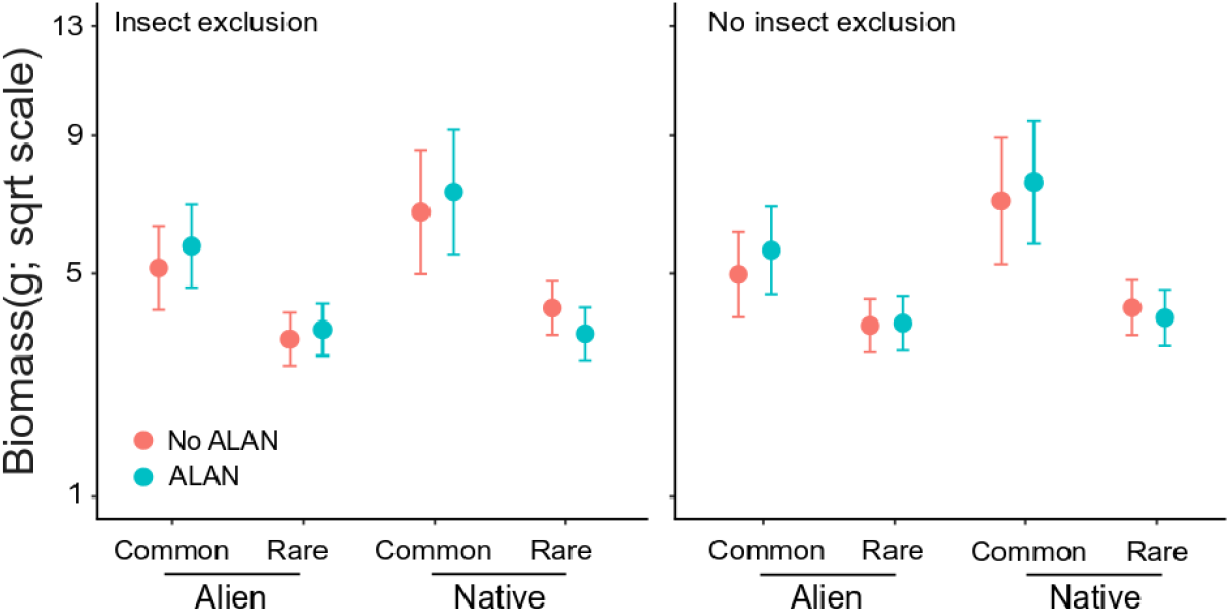
Modelled mean values of biomass per pot for different ALAN treatments (no ALAN *vs* ALAN), species commonness (common *vs* rare) and species origin (native *vs* alien) in the cages with insect exclusion and the cages without insect exclusion. Error bars represent standard errors.

**Figure 3.**
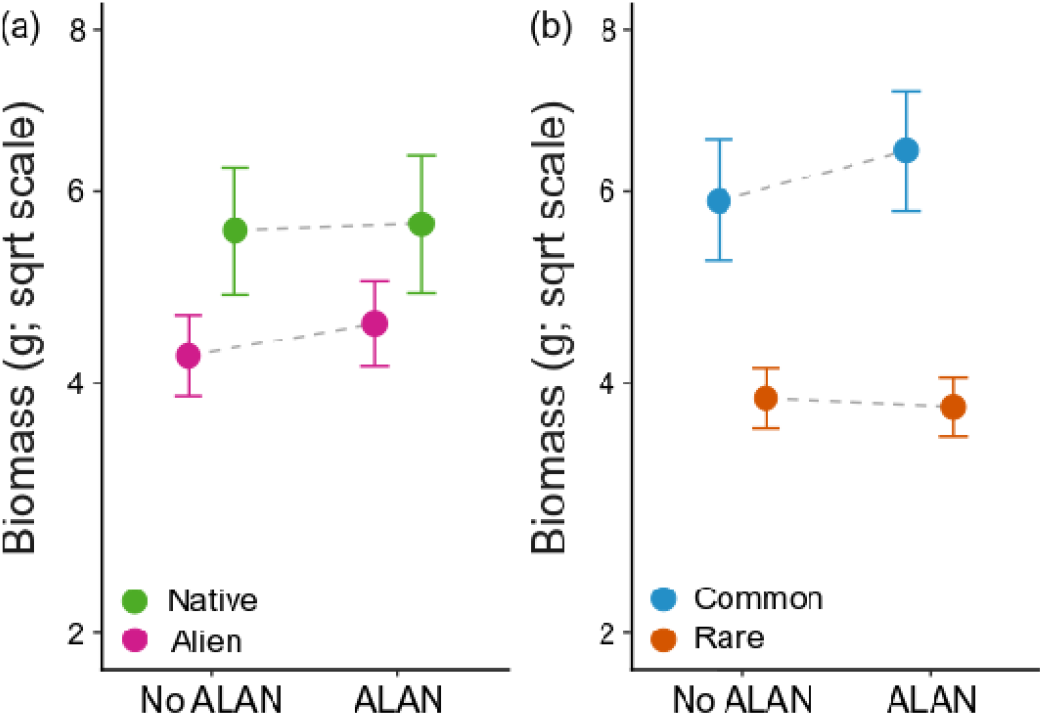
Modelled mean values of biomass per pot for different ALAN treatments (no ALAN *vs* ALAN) and species origin (native *vs* alien) (a) and for different ALAN treatments (no ALAN *vs* ALAN) and species commonness (common *vs* rare) (b). Error bars represent standard errors.

## Discussion

Our study tested how ALAN interacts with the presence of insects to affect the performance of common alien, rare alien, common native and rare native species. We found that the common species, irrespective of whether they are alien or native, had a significantly higher performance than rare ones, and that this difference was even amplified by ALAN. Furthermore, alien species tended to show a slightly more positive response to ALAN than native species (marginally significant origin × ALAN interaction), indicating that ALAN might promote alien plant invasion.

It has been suggested that common alien species and common native species are both successful, and thus should share similar attributes ^18,20,45,46^. Furthermore, many common native species have successfully naturalized elsewhere (e.g. Ref. ^47^). Indeed, our multispecies experiment showed that common plant species generally produced more aboveground biomass than rare species, regardless of whether they are alien or native. It is also worth noting that the aboveground biomass production of common plant species increased to a greater degree with ALAN than it did for rare species. This most likely reflects that even the faint light emitted by street-lamps is bright enough for plants to do photosynthesis, partly or entirely compensating dark respiration ^15^, and that common species often have greater ability than rare ones to capture light ^25,26^. So, our results corroborate those of Dawson et al. ^18^, who found that common species benefit more from additional resources (i.e. nutrients in the case of Ref. ^18^) than rare species do.

The present findings, however, contrast with the results of Speißer et al. ^15^, who found that the less widely naturalized species tended to increase their biomass more strongly in response to ALAN than the widely naturalized species did. A possible explanation for this could be that the two studies differed in the number of dimensions of commonness that they used ^48,49^. While Speißer et al. ^15^ classified species as common or rare based on their grid-cell occurrence frequency (in Germany) only, we here used local abundance as an additional, more restrictive, criterion. Therefore, future studies assessing how ALAN affects performance of common and rare plants should test how ALAN affects the different dimensions of commonness. Nevertheless, our present study highlights that the ongoing increase in ALAN caused by urbanization may trigger the so-called “Matthew Effect”, i.e. that common species might become even more common.

ALAN is particularly common along roadsides ^6,8^, which are also important invasion corridors for alien plants ^50,51^. Indeed it was recently shown that the presence of an invasive grass species was positively associated with the presence of streetlights ^17^. Our experimental study provides further evidence that alien plant invasion might be facilitated by ALAN, as the alien plants tended to respond more positively to ALAN than native plants. The marginally greater biomass response to ALAN for alien plants may have been due to their higher resource-use efficiency ^52,53^ and greater phenotypic plasticity in response to light ^54-56^. Although plants in our study did not grow under interspecific competition, our results suggest that the slight increase in biomass of alien plants in response to ALAN might help them when they compete with native plants.

In addition to direct effects, we expected that ALAN might also have indirect effects on alien plant invasion via other trophic levels. This is because alien plants often attract different numbers and types of insects than native plants do ^35,36,38^, and ALAN could also modify plant-insect interactions ^7,9,31,32^. Our study, however, found no evidence that insect exclusion mediated the effects of ALAN on alien and native plants. Actually, the main effect of insect exclusion was also not significant, although we caught many more flying insects in the open cages than in the closed cages. Most of the insects, however, were Chironomidae (Table S2), and, although some of them might feed on nectar, they do not damage plant tissue and their role in pollination is not clear ^57-59^. Thus the absence of large numbers of herbivores in our study location most likely explains why we did not find an effect of insect exclusion on biomass production.

In conclusion, our study showed that common alien and native plant species benefit more from ALAN than rare plant species. Such differences might lead to shifts in plant diversity and vegetation composition in grasslands with further increases of ALAN. In addition, the slightly more positive effect of ALAN on the alien plant species than on the native ones indicates that increased ALAN might also further increase the invasion risk of the alien species.

## Acknowledgements

We thank Beate Rüter, Heinz Vahlenkamp, Maximilian Fuchs, Otmar Ficht, Sarah Berg and Vanessa Pasqualetto for their help. YL was supported by the Chinese Academy of Sciences (project Y9B7041001), and the Zukunftskolleg of the University of Konstanz (Independent Research Grant).

## Author contributions

YL conceived the idea and designed the experiment with inputs from MvK and BS. BS performed the experiment. YL analyzed the data and wrote the first draft of the manuscript, with further inputs from BS, EK, and MvK.

## Data accessibility

Should the manuscript be accepted, the data supporting the results will be archived in Dryad and the data DOI will be included at the end of the article.

## Supporting information

**Table S1.** Details of the study species used in the experiment.

| Family | Species | Origin | Commonness | Number of grid cells* | Life History | Origin | Sowing date |
| --- | --- | --- | --- | --- | --- | --- | --- |
| Amaranthaceae | *Amaranthus graecizans* L. | Alien | Rare | 76 | Annual | Botanical Garden Regensburg | 13.05.2019 |
| Amaranthaceae | *Amaranthus retroflexus* L. | Alien | Common | 2084 | Annual | B and T World Seeds SARL | 20.05.2019 |
| Amaranthaceae | *Chenopodium rubrum* L. | Native | Common | 1919 | Annual | Botanical Garden Konstanz | 20.05.2019 |
| Asteraceae | *Centaurea calcitrapa* L. | Alien | Rare | 170 | Biennial | Botanical Garden Konstanz | 20.05.2019 |
| Asteraceae | *Gnaphalium uliginosum* L. | Native | Common | 2845 | Annual | Botanical Garden Konstanz | 20.05.2019 |
| Asteraceae | *Senecio vernalis* Waldst. & Kit | Alien | Common | 2031 | Annual | Botanical Garden Konstanz | 20.05.2019 |
| Brassicaceae | *Cardamine amara* L. | Native | Common | 2628 | Perennial | Botanical Garden Konstanz | 20.05.2019 |
| Brassicaceae | *Lepidium graminifolium* L. | Native | Rare | 86 | Perennial | Botanical Garden Konstanz | 20.05.2019 |
| Brassicaceae | *Lepidium heterophyllum* Benth. | Alien | Rare | 98 | Perennial | Botanical Garden Konstanz | 20.05.2019 |
| Fabaceae | *Galega officinalis* L. | Alien | Rare | 259 | Perennial | Botanical Garden Jibou | 20.05.2019 |
| Fabaceae | *Trifolium ochroleucon* Huds. | Native | Rare | 300 | Perennial | Botanical Garden Konstanz | 20.05.2019 |
| Fabaceae | *Vicia cracca* L. s. str. | Native | Common | 2918 | Perennial | Revierberatung Wolmersdorf | 20.05.2019 |
| Fabaceae | *Vicia villosa* Roth s. I. | Alien | Common | 1993 | Annual | Revierberatung Wolmersdorf | 20.05.2019 |
| Lamiaceae | *Hyssopus officinalis* L. | Alien | Rare | 199 | Perennial | Botanical Garden Konstanz | 20.05.2019 |
| Lamiaceae | *Leonurus marrubiastrum* L. | Native | Rare | 191 | Annual | Botanical Garden Konstanz | 20.05.2019 |
| Lamiaceae | *Salvia verticillata* L. | Alien | Common | 977 | Perennial | Rieger-Hofmann GmbH | 13.05.2019 |
| Poaceae | *Bromus hordeaceus* L. | Native | Common | 2928 | Annual | Rieger-Hofmann GmbH | 20.05.2019 |
| Poaceae | *Bromus japonicus* Thunb. | Alien | Rare | 355 | Biennial | Botanical Garden Konstanz | 20.05.2019 |
| Poaceae | *Eragrostis minor* Host | Alien | Common | 1448 | Annual | Natural community (Konstanz) | 20.05.2019 |
| Poaceae | *Parapholis strigosa* (Dumort.) C.E. Hubb. | Native | Rare | 84 | Annual | Botanical Garden Paris | 20.05.2019 |

**Table S2.** The numbers of insects caught per 4 cm^2^ of insect trap in each cage. We had two 5 cm × 12 cm sticky, yellow traps per cage, one at plant level and one below the lamp shades. On each stick trap, the numbers of insects were counted in a 1 cm × 2 cm area at the bottom right corner of the traps.

| Cage ID | ALAN treatment | Insect exclusion treatment | Number of insects |  |  |
| --- | --- | --- | --- | --- | --- |
|  |  |  | Chironomidae | Unidentified | Total |
| 1 | ALAN | No insect exclusion | 20 | 9 | 29 |
| 2 | No ALAN | Insect exclusion | 0 | 0 | 0 |
| 3 | ALAN | No insect exclusion | 11 | 7 | 18 |
| 4 | No ALAN | Insect exclusion | 0 | 1 | 1 |
| 5 | ALAN | No insect exclusion | 39 | 3 | 42 |
| 6 | No ALAN | Insect exclusion | 0 | 1 | 1 |
| 7 | No ALAN | No insect exclusion | 0 | 3 | 3 |
| 8 | No ALAN | No insect exclusion | 0 | 2 | 2 |
| 9 | ALAN | No insect exclusion | 22 | 1 | 23 |
| 10 | No ALAN | Insect exclusion | 0 | 0 | 0 |
| 11 | ALAN | Insect exclusion | 2 | 2 | 4 |
| 12 | ALAN | Insect exclusion | 4 | 0 | 4 |
| 13 | ALAN | No insect exclusion | 15 | 14 | 29 |
| 14 | No ALAN | No insect exclusion | 2 | 2 | 4 |
| 15 | No ALAN | No insect exclusion | 0 | 0 | 0 |
| 16 | No ALAN | Insect exclusion | 0 | 0 | 0 |
| 17 | ALAN | Insect exclusion | 6 | 4 | 10 |
| 18 | ALAN | Insect exclusion | 7 | 2 | 9 |
| 19 | ALAN | Insect exclusion | 9 | 3 | 12 |
| 20 | No ALAN | No insect exclusion | 0 | 2 | 2 |

**Table S3.** Standard deviation and multiplication factors for individual species random effects for the model shown in Table 1 of the main manuscript. The standard deviation given refers to the first species (i.e., LH). For each other species, the standard deviation should be multiplied by the respective multiplication factor. The names of the species in the table are abbreviated using the first letter of the genus and species epithet (for species identity, see Table S1).

| Metric | Species Standard<br>Deviation | Multiplication factor standard deviation |  |  |  |  |  |  |  |  |  |  |  |  |  |  |  |  |  |  |  |
| --- | --- | --- | --- | --- | --- | --- | --- | --- | --- | --- | --- | --- | --- | --- | --- | --- | --- | --- | --- | --- | --- |
|  |  | LH | VV | TO | CA | CC | LG | SV | LM | GO | HO | BH | AG | PS | AR | BJ | SV | EM | CR | VC | GU |
| Aboveground<br>biomass | 0.469 | 1.000 | 1.625 | 0.881 | 0.732 | 0.637 | 0.572 | 0.710 | 0.587 | 1.035 | 0.935 | 0.615 | 0.627 | 0.795 | 0.980 | 0.566 | 1.002 | 1.096 | 1.043 | 1.071 | 0.943 |

**Figure S1.**
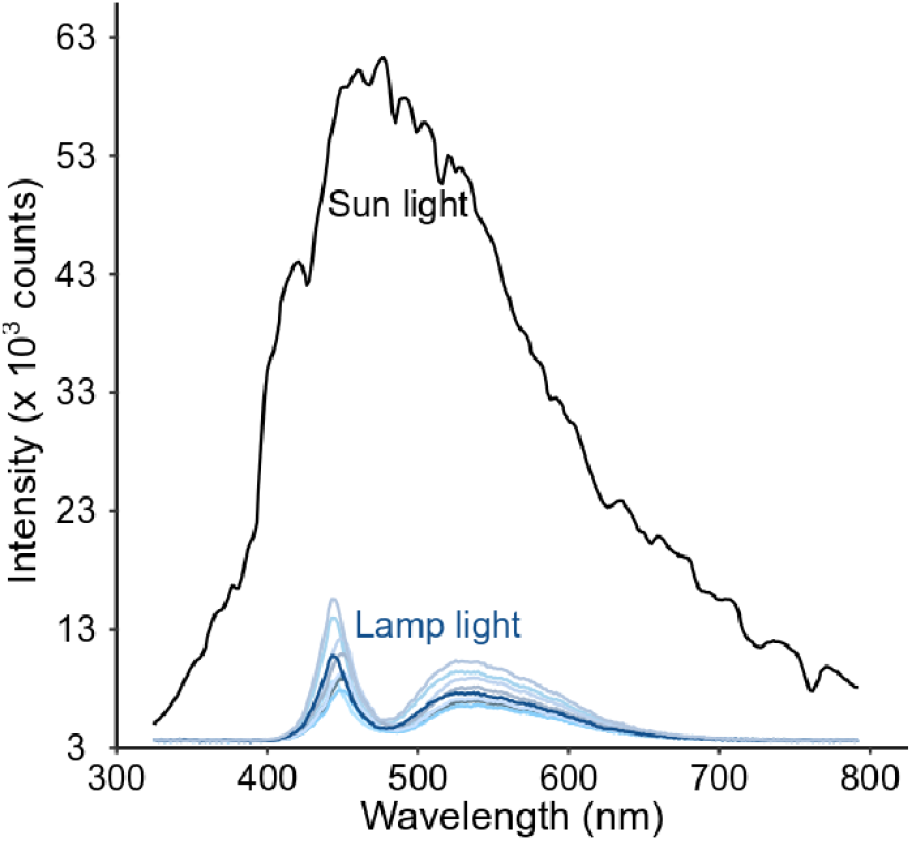
Spectral composition of sunlight (black line), artificial light emitted by the ten LED spotlights (blue lines). Photosynthetically active radiation is in the range from 400 to 700nm.

**Figure S2.**
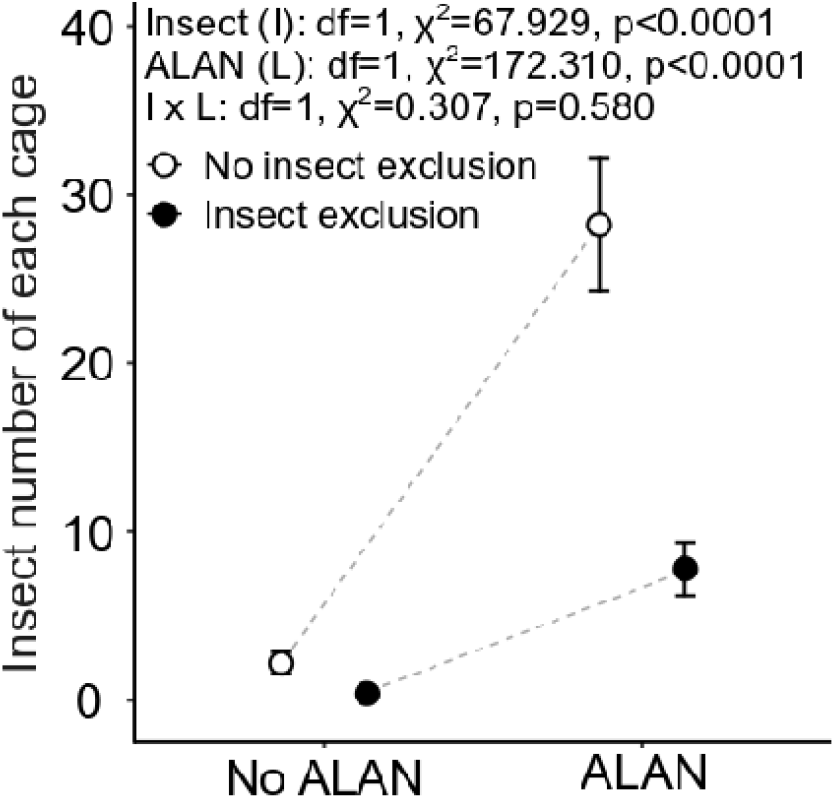
Mean values of insect numbers caught per cage in the different ALAN treatments (no ALAN *vs* ALAN) and different insect-exclusion treatments (no exclusion *vs* exclusion). Error bars represent standard errors.

